# Fluorescent non-canonical amino acid as a site-specific conformational probe of prion formation

**DOI:** 10.64898/2026.04.21.719968

**Authors:** Jessica de Alcantara Ferreira, Daniel J. Walsh, Evelyn Turnbaugh, Jeremy H. Mills, Surachai Supattapone

**Affiliations:** Department of Biochemistry, Geisel School of Medicine at Dartmouth, Hanover, New Hampshire 03755, USA; Department of Medicine, Geisel School of Medicine at Dartmouth, Hanover, New Hampshire 03755, USA; School of Molecular Sciences and The Biodesign Center for Molecular Design and Biomimetics, Arizona State University, Tempe, AZ 85287, USA

**Keywords:** prion, non-canonical amino acid, conformation, 7-hydroxycoumarin, amyloid

## Abstract

The pathogenic conversion of the cellular prion protein (PrP^C^) into the β-sheet-rich isoform PrP^Sc^ is the pivotal pathogenic event in prion disease, yet the molecular steps that govern this structural transition remain elusive. In this study, we introduce a new approach to monitor site-specific conformational transitions that occur during infectious prion formation. The method relies on genetically encoded substitution of a fluorescent, environmentally sensitive non-canonical amino acid, L-(7-hydroxycoumarin-4-yl)ethylglycine (7-HCAA), into recombinant PrP substate molecules, allowing real-time monitoring of structural changes in high-efficiency *in vitro* PrP^Sc^ conversion reactions. As proof of principle, we show that the W99 7-HCAA recPrP substate efficiently propagates two different PrP^Sc^ conformers (infectious cofactor PrP^Sc^ and non-infectious protein-only PrP^Sc^). Bioassays in knock-in mice expressing bank vole (BV) PrP confirm that W99 7-HCAA cofactor PrP^Sc^ produced by serial propagation is infectious, causing scrapie with an incubation period and neuropathological profile like those induced by wild-type cofactor PrP^Sc^. Marked differences in fluorescence intensity were observed between native, misfolded, and denatured states of W99 7-HCAA PrP, confirming that 7-HCAA reports on local changes in PrP conformation. Together, these findings establish 7-HCAA as a site-specific and sensitive probe of local PrP conformation. Moreover, the results suggest a new and broadly applicable strategy for studying conformational dynamics in amyloid-forming proteins.

## Introduction

Prions are unconventional infectious agents that cause fatal neurological diseases, including Creutzfeldt-Jakob disease (CJD), bovine spongiform encephalopathy (BSE), and scrapie. Prion diseases can be either spontaneous (sporadic), genetic, or acquired via infection*(1, 2)*. The causative event in prion disease is the misfolding of the host-encoded prion protein (PrP^C^) into misfolded, insoluble, and protease-resistant conformers (collectively termed PrP^Sc^) through a self-propagating mechanism*(3, 4)*.

Cryogenic electron microscopy(cryo-EM) studies show that PrP^Sc^ molecules adopt parallel in-register β-sheet (PIRBS) structures within amyloid fibrils*(5-9)*. Although the structures of PrP^C^*(10)* and several PrP^Sc^ conformers have been solved, the structural mechanism underlying PrP^C^ to PrP^Sc^ conversion remains unknown. Molecular dynamics simulations and biophysical studies suggest that an early and critical event is the destabilization of helix 1, the most labile element of PrP^C^, followed by its separation from the helix 2/helix 3 core to expose hydrophobic surfaces that nucleate β-rich oligomers*(11, 12)*. Although these *in silico* studies provide useful hypotheses to test, there are currently no experimental techniques available to study the dynamic structural transition from PrP^C^ to PrP^Sc^.

It has been technically challenging to monitor the conversion of PrP^C^ to PrP^Sc^ in part because it has not previously been possible to specifically label PrP^C^ with a conformationally sensitive probe without disrupting its structure and its ability to convert to infectious PrP^Sc^. Current labeling methods include genetic insertion of large tags or conjugation of chemically reactive probes*(13-15)*, but these substantial manipulations usually destabilize the modified PrP^C^ structure and prevent it from converting into PrP^Sc^. In contrast, small non-canonical amino acids (ncAA) with diverse chemical properties can be inserted into specified sites within recombinant proteins with minimal perturbation*(16-20)*

Here, we use 7-hydroxycoumarin-4-yl ethylglycine (7-HCAA) as a probe to monitor PrP misfolding during the process of prion formation *in vitro*. 7-HCAA is a fluorescent ncAA whose fluorescence properties (e.g., maximal emission wavelength and/or emission intensity) can be altered by protein environments. These properties have led to the use of 7-HCAA in previous studies of protein-protein and protein-ligand interaction*(21-26)*. Expressing proteins with genetically encoded 7-HCAA requires an orthogonal tRNA/aminoacyl-tRNA synthetase (aaRS) pair that selectively recognizes 7-HCAA and charges only its specific orthogonal tRNA *(27, 28)*. The orthogonal tRNA is engineered with a CUA anticodon to decode the amber stop codon (UAG). Incorporation of 7-HCAA thus occurs by suppressing an in-frame amber codon introduced at the desired position by site-directed mutagenesis*(29-31)*. Here we combined this method of producing specific 7-HCAA-substituted PrP substrates with a high efficiency infectious prion conversion system*(32-35)* to create a novel platform to study the dynamic process of PrP^Sc^ formation.

## Materials and Methods

### Animal husbandry

The Guide for the Care and Use of Laboratory Animals of the National Research Council was strictly followed for animal experiments. Mice breeding and euthanasia was conducted in accordance with protocols 00002131 and 00002140 as reviewed and approved by Dartmouth College’s Institutional Animal Care and Use Committee, operating under the regulations/guidelines of the NIH Office of Laboratory Animal Welfare (assurance number A3259-01).

## Cloning of PrP construct containing 7-HCAA

The insertion of the non-canonical amino acid 7-HCAA into the M109 BV PrP 23-23 sequence (UniProt ID: Q8VHV5_MYOGA) was achieved through the amber code suppression method established by Schultz *et al*.*(16, 36)*. PrP sequences containing 7-HCAA substitution were generated by Gibson assembly. pET-22b(+) (Sigma-Aldrich, St. Louis, MO) was digested with NdeI and HIndIII restriction enzymes (New England Biolabs, Ipswich, MA), gel-purified, and assembled together with a G-block encoding M109 BV PrP 23-23 (codon-optimized for bacteria) with appropriate flanking regions and amber substitution (IDT, Newark, NJ) using the Gibson Assembly^®^ Master Mix (New England Biolabs, Ipswich, MA), following manufacturer’s instructions. The backbone sequence for G-blocks (without amber substitution) is:

5’-

TTG TTT AAC TTT AAG AAG GAG ATA TAC ATA TGA AGA AGC GTC CTA AAC CTG GGG GGT GGA ACA CCG GTG GTA GCC GTT ATC CTG GGC AAG GCA GCC CAG GTG GAA ACC GTT ACC CTC CTC AAG GTG GCG GGA CTT GGG GAC AGC CCC ACG GTG GAG GCT GGG GAC AGC CTC ATG GAG GGG GTT GGG GCC AAC CGC ATG GCG GTG GCT GGG GTC AAC CTC ATG GAG GCG GGT GGG GAC AGG GGG GAG GAA CAC ATA ATC AAT GGA ATA AAC CCA GTA AGC CTA AAA CAA ACA TGA AGC ACG TAG CCG GTG CGG CCG CAG CCG GAG CAG TCG TGG GGG GAT TGG GCG GAT ATA TGC TTG GAT CGG CAA TGA GTC GCC CCA TGA TCC ATT TCG GAA ATG ACT GGG AAG ATC GCT ATT ATC GCG AAA ACA TGA ACC GCT ATC CGA ACC AGG TTT ACT ATC GCC CGG TTG ATC AAT ACA ACA ATC AAA ACA ACT TTG TAC ATG ATT GCG TTA ATA TTA CAA TCA AAC AGC ACA CAG TTA CCA CAA CTA CTA AAG GCG AGA ATT TTA CTG AGA CGG ACG TCA AAA TGA TGG AGC GTG TAG TTG AGC AGA TGT GTG TCA CAC AAT ATC AGA AGG AAT CGC AAG CCT ATT ACG AAG GCC GCT CGT CCT GAT AAA AGC TTG CGG CCG CAC TCG AGC ACC ACC ACC A-3’

The resulting plasmid was transformed into DH5α cells (Invitrogen, Waltham, MA), purified using a miniprep kit (Qiagen, Hilden, Germany), and verified by DNA sequencing.

### Expression and purification of recPrP containing the non-canonical amino acid 7-HCAA

*E. coli* BL21 derivative B-95.ΔAΔ*fabR* cells (kindly provided by Kensaku Sakamoto, RIKEN Systems and Structural Biology Center, Yokohama, Japan) were co-transformed with pET22b plasmid containing an ampicillin resistance marker and genes encoding the mutant PrP, and pEVOL-CouRS plasmid*(23)* (kindly provided by Peter Schultz, Scripps Research, La Jolla, California, USA, and Jeremy Mills, Arizona State University, Arizona, USA) containing chloramphenicol resistance marker, two copies of the evolved CouRS synthetase, and an engineered tRNA that is recognized and charged with 7-HCAA by the evolved CouRS and decodes the UAG codon. Double transformants were selected on LB agar plates. A single *E. coli* colony was used to inoculate 3.0 mL of 2xYT medium in a sterile round-bottom culture tube. The culture was incubated overnight at 37°C with agitation at 230 rpm. Cells were harvested by centrifugation at 4200 × *g* for 10 min and resuspended in 3 mL of fresh 2×YT medium lacking antibiotics. This suspension was used to inoculate 250 mL of 2×YT, also lacking antibiotics, in a baffled flask.

Cultures were grown at 37°C with shaking until OD_600_ reached ∼3.0. At this point, expression of the orthogonal aminoacyl-tRNA synthetase (CouRS) was induced by the addition of 0.2% (w/v) L-arabinose and 1 mM of the non-canonical amino acid 7-HCAA (*M*_r_ ∼ 263.25 kDa, catalog # 4059541, Bachem, Torrance, CA). Cultures were incubated for an additional hour under the same conditions. Protein expression was then induced by adding isopropyl-β-D-thiogalactoside (IPTG) (catalog # BP1755-1, Fisher, Santa Clara, CA) to a final concentration of 1 mM, followed by incubation at 30°C with shaking at 230 rpm for 16 hr. Bacteria were pelleted at 4,000 x g for 25 min. Unless otherwise noted, all procedures were carried out in the presence of 34 µg/mL chloramphenicol and 50 µg/mL ampicillin.

The resulting protein was purified following the method published by Makarava *et al*., which ultimately includes metal ion affinity chromatography, size-exclusion chromatography (for desalting), and reverse-phase chromatography*(37-39)*. Lyophilized protein was resuspended in water and adjusted to a concentration of 0.12 mg/mL based on absorbance at A280 before use.

### *In vitro* propagation of PrP^Sc^

Substrate cocktails for recPrP conversion reactions were prepared as previously described by Noble *et al*.*(38)*. Briefly, substrate cocktails contain 6 μg/mL recombinant BV PrP (residues 23– 231, either wild-type (WT) or W99 7-HCAA recPrP) in conversion buffer consisting of 20 mM Tris, 135 mM NaCl, 5 mM EDTA, 0.15% (v/v) Triton X-100, pH 7.4, and optional lipid cofactor (from 10-20%). *In vitro* conversion reactions were initiated by combining 100 µL of 6 µg/mL PrP^Sc^ (in reaction buffer) with 400 µL of substrate shaking cocktail, followed by incubation with shaking for 72 hr at 37°C. For serial propagation, 100 µL of the newly generated PrP^Sc^ was transferred into a fresh 400 µL substrate cocktail to initiate the subsequent round of amplification. W99 7-HCAA cofactor PrP^Sc^ was generated by seeding a cofactor-containing W99 7-HCAA recPrP substrate cocktail with WT cofactor PrP^Sc^. To produce W99 7-HCAA protein-only PrP^Sc^, the W99 7-HCAA PrP^C^ substrate mixture was initially seeded with WT cofactor PrP^Sc^, followed by gradual elimination of the cofactor over ∼30 serial propagation rounds.

Shaking was performed in a custom-built shaker with an orbital diameter of 8 mm at approximately 1,200 rpm. To prepare samples for inoculation, samples initially seeded with either cofactor PrP^Sc^ or protein-only PrP^Sc^ samples were serially propagated for 26 rounds to produce inocula.

### Prion inoculation and neuropathology

Intracerebral inoculation and diagnosis of prion disease were performed as described previously*(40)* with the following modifications. Reaction mixtures containing PrP^Sc^ were diluted 1:10 in PBS 1% (w/v) bovine serum albumin to be used as inoculum. 30 μL of the inoculum was injected into kiBV M109 PrP mice (breeding pairs kindly provided by Joel C. Watts, University of Toronto, Ontario, Canada) between 4–6 weeks of age.

Brains were removed rapidly at the time of sacrifice using new, sterile-packaged dissection instruments. They were immersion-fixed in 10% buffered formalin for 2–30 days, cut into ∼3 mm-thick sagittal sections, and placed in a tissue-processing cassette. Cassettes were treated with 88% formic acid for 1 hr and then stored in PBS. The tissue was processed for paraffin embedding, and representative slides were stained with hematoxylin and eosin (H&E) (performed at Dartmouth-Hitchcock Medical Center, Department of Pathology & Laboratory Medicine).

### Proteinase K digestion of PrP^Sc^

PrP^Sc^ samples produced *in vitro* were treated with 20 μg/mL proteinase K (PK) (catalog # 3115828001, Millipore Sigma) at 37°C with shaking at 750 rpm for 30 min. Brain homogenates (10% (w/v) in PBS) from experimentally infected brains were digested in a reaction containing 100 μg/mL PK, 1%(v/v) Triton at 37°C with shaking at 800 rpm for 30 min. All digestion reactions were quenched by the addition of 4 mM PMSF (catalog #P7626, Millipore Sigma).

### PrP^Sc^ detection

PrP^Sc^ detection was performed both through visualization of fluorescent gel and Western blot. To assess PrP^Sc^ electrophoretic mobility after protease digestion, samples were denatured by boiling for 15 min in Laemmli SDS sample buffer (catalog # SAB03-01, Bioland Scientific, Paramount, CA). SDS–polyacrylamide gel electrophoresis (SDS-PAGE) was performed using 1.5-mm-thick, 12% polyacrylamide gels prepared with a 29:1 acrylamide to bisacrylamide ratio. Following electrophoresis, the gel was imaged for fluorescence using the Azure 600 system (catalog # AZI600-01, Azure Biosystems, Dublin, CA), with excitation wavelength set for 365 nm and emission at 513 nm.

For Western blot detection, proteins were transferred to methanol-activated Millipore Immobilon-P polyvinylidene difluoride (PVDF) membrane (Millipore, Billerica, MA) using a Trans-Blot SD semi-dry transfer system (catalog # 170-3940, Bio-Rad, Hercules, CA) at a constant current of 2.5 mA/cm^2^ for 50 minutes. To detect PrP^Sc^, membranes were blocked with nonfat dry milk in TBST buffer (10 mM Tris, pH 7.1, 150 mM NaCl, 0.1% Tween 20) for 1 hr, then washed 3 times in TBST for 10 minutes. Following that, the membrane was incubated for 2 hr at room temperature with the 27/33 monoclonal antibody diluted 1:25,000 (final concentration: 0.12 μg/mL). After three 10-minute washes with TBST, membranes were incubated for 1 hr with a horseradish peroxidase-conjugated anti-mouse IgG secondary antibody (GE Healthcare) diluted 1:5,000 in TBST. Incubation with the antibody was followed by three 10-minute washes in TBST. Signal detection was performed using West Pico Plus chemiluminescent substrate (catalog #22169, Thermo Fisher Scientific), and images were captured with the same Azure system. Molecular weight estimations were based on pre-stained protein markers (Fermentas, Hanover, MD).

### Assessment of 7-HCAA fluorescence levels in solution samples

Samples were split into two equal halves, unless otherwise specified, and fluorescence was simultaneously measured under (1) denaturing conditions using 6 M guanidine HCl, 20 mM Tris, pH 7.4 and boiling samples at 60°C for 30 minutes to quantify total fluorescence, and (2) non-denaturing conditions using control buffer (20 mM Tris, pH 7.4) to assess fluorescence in the native or fibrillar state. A SpectraMax^®^ iD5 (Molecular Devices) plate reader was used to record fluorescence spectra from 400 to 500 nm at 10 nm intervals, unless otherwise specified. Measurements were performed in biological triplicate, with excitation at 360 nm, to acquire a fluorescence emission curve. For fluorescence endpoint measurements, a filter was used to select 360 nm excitation and 460 nm emission wavelengths. Background fluorescence was subtracted from all sample measurements prior to analysis. Baseline values were obtained from buffer-only controls measured under identical conditions. The fluorescence intensity data were plotted and analyzed using GraphPad Prism.

### Statistical Analysis

All statistical analyses were carried out using GraphPad Prism (version 10.6.1). Data are presented as mean ± standard error of the mean (SEM). Two-tailed Welch’s t-test was used to compare means between two independent groups. For comparisons across three or more groups, ordinary one-way ANOVA was performed, followed by Tukey’s post-hoc test for multiple comparisons. Statistical significance was defined as p < 0.05.

## Acknowledgements

This study was funded by the National Institute for Neurological Diseases and Stroke (1R37NS125431, R01NS117276 and R01NS118796 to S.S.) and the National Institutes of Health (P20-GM113132 to Dean Madden).

We would like to thank Kensaku Sakamoto (RIKEN Systems and Structural Biology Center, Yokohama, Japan) for kindly providing *E. coli* BL21 derivative B-95.ΔAΔ*fabR* cells.

## Results

### W99 7-HCAA recPrP substrate can be efficiently converted into PrP^Sc^

We first expressed a variety of mutant recPrP molecules in which individual aromatic residues were substituted with 7-HCAA in *E. coli*. Analysis of crude bacterial lysates by fluorescence gel imaging showed that most of the mutant 7-HCAA recPrP molecules were successfully expressed as fluorescent, full-length proteins **(Supplemental Figure 1)**.

**Fig 1:**
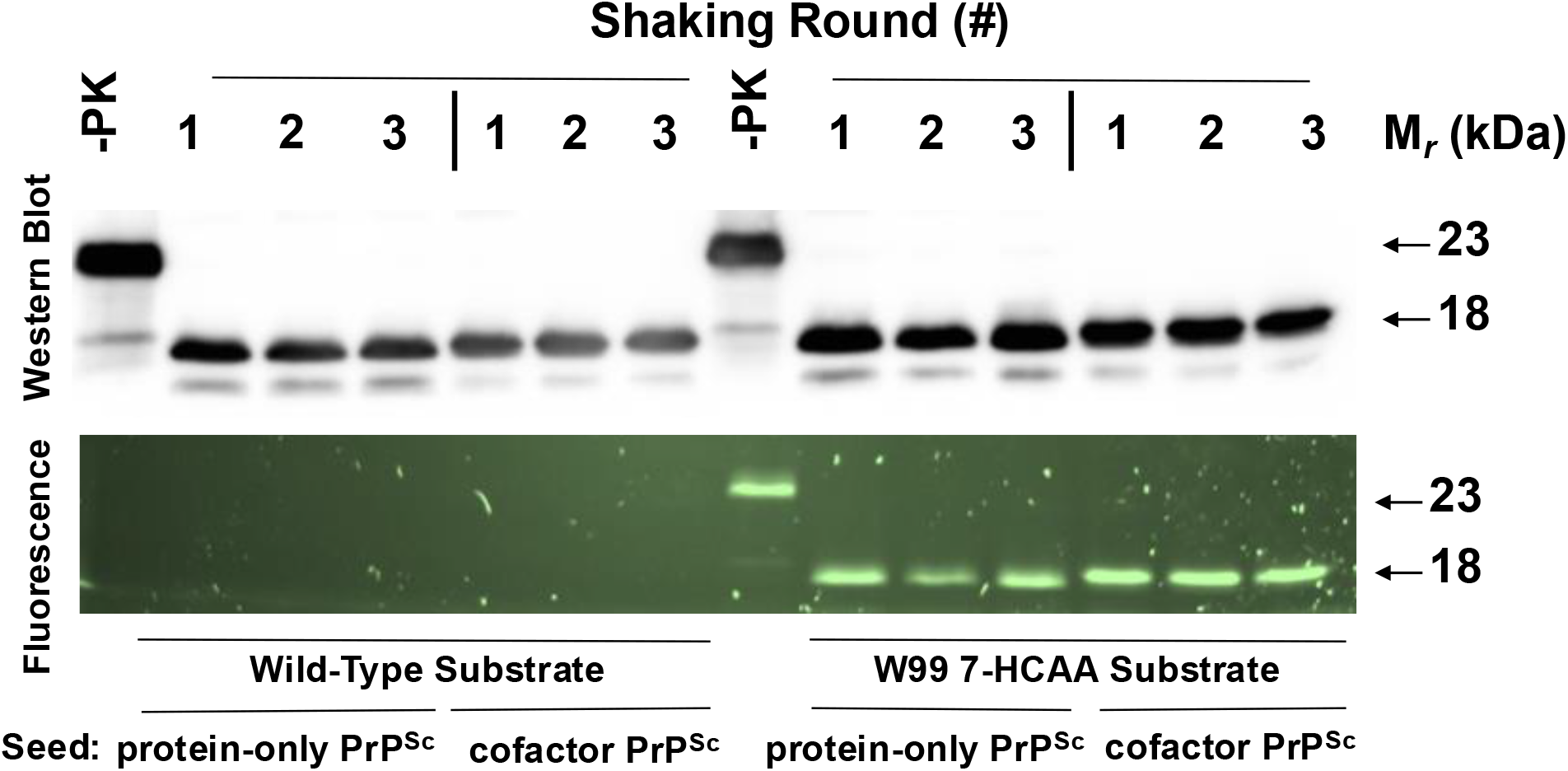
Serial propagation of two PrP^Sc^ conformers using WT and W99 7-HCAA recPrP substrates. Western blot probed with anti-PrP mAb 27/33 (Top panel) and corresponding fluorescent gel (Bottom panel) showing three rounds of serially propagated reactions using either wild-type (WT) or W99 7-HCAA substrate, as indicated, and initially seeded with either protein-only PrP^Sc^ or cofactor PrP^Sc^, as indicated. -PK = sample not subjected to proteinase K digestion; all other samples were proteolyzed.

We next tested whether a representative mutant PrP molecule (W99 7-HCAA recPrP) could function as a substrate for a high-efficiency *in vitro* PrP conversion system that can propagate either a highly infectious PrP^Sc^ conformer (termed cofactor PrP^Sc^) or a non-infectious PrP^Sc^ conformer (termed protein-only PrP^Sc^), depending on the initial seed and addition of phospholipid cofactor to the reaction mixture*(33, 34, 41)*. We purified W99 7-HCAA PrP and used it as a substrate in 3-round serial propagation reactions initially seeded with either cofactor PrP^Sc^ or protein-only PrP^Sc^. Western blot analysis shows that cofactor PrP^Sc^-seeded reactions using W99 7-HCAA recPrP substrate generated self-propagating PrP^Sc^ molecules **(Figure 1, top, lanes 12-14)**. On Western blot, the PK-resistant core for W99 7-HCAA cofactor PrP^Sc^ had a molecular weight of ∼18 kDa, similar to the molecular weight of wild-type (WT) cofactor PrP^Sc^ **(Figure 1, top, compare lanes 5-7 with 12-14)**. The corresponding fluorescent gel shows fluorescence when excited at 365 nm for samples with W99-7HCAA recPrP as substrate, confirming 7-HCAA incorporation **(Figure 1, bottom, lanes 12-14)**, whereas samples using WT recPrP substrate were not fluorescent **(Figure 1, bottom, lanes 5-7)**. W99 7-HCAA recPrP substrate also converted efficiently into self-propagating, fluorescent PrP^Sc^ molecules in protein-only PrP^Sc^-seeded reactions **(Figure 1 top and bottom, lanes 9-11)**. The PK-resistant core of W99 7-HCAA protein-only PrP^Sc^ displayed a molecular weight of ∼17 kDa, similar to that of WT protein-only PrP^Sc^*(35, 37)* **(Figure 1, top, compare lanes 2-4 with 9-11)**. Taken together, the results show that W99 7-HCAA recPrP can be efficiently converted into and propagated as either W99 7-HCAA cofactor PrP^Sc^ or W99 7-HCAA protein-only PrP^Sc^.

### Cofactor W99 7-HCAA PrP^Sc^ seeds are infectious *in vivo*

To examine if the W99 7-HCAA cofactor PrP^Sc^ produced in our serial shaking reaction is infectious, we performed 26 serial propagation cycles to ensure that all the WT cofactor PrP^Sc^ molecules in the initial seed were diluted beyond Avogadro’s limit. We then inoculated kiBV M109 PrP mice*(42)* with the round 26 product. We also included unconverted W99 7-HCAA cofactor recPrP substrate cocktail and W99 7-HCAA protein-only PrP^Sc^ as negative controls, and WT cofactor PrP^Sc^ as a positive control.

Mice inoculated with W99 7-HCAA cofactor PrP^Sc^ developed scrapie ∼ 180 days after inoculation, showing hunched posture, ataxia, and circling behavior **(Figure 2A, red squares)**. Similarly, mice inoculated with wild-type (WT) cofactor PrP^Sc^ developed scrapie ∼200 days after inoculation **(Figure 2A, blue circles)**. In contrast, mice inoculated with unconverted W99 7-HCAA cofactor recPrP substrate cocktail or with W99 7-HCAA protein-only PrP^Sc^ survived for at least 400-600 days **(Figure 2A, see green diamond and black triangle)**. Western blot analysis revealed the accumulation of PK-resistant PrP^Sc^ in the brains of symptomatic mice inoculated with cofactor W99 7-HCAA PrP^Sc^, with a similar pattern to the PK-resistant core of PrP^Sc^ in the brains of animals inoculated with WT cofactor PrP^Sc^ **(Figure 2B, compare lanes 5 and 6 to lanes 7 and 8)**. The brains of the mice inoculated with W99 7-HCAA cofactor PrP^Sc^ show abundant vacuolation, whereas brains inoculated with unconverted substrate or WT protein-only PrP^Sc^ samples were histologically normal **(Figure 3, compare panels A and B *versus* panel C)**. The distribution of vacuolation in different brain regions was also similar between mice inoculated with W99 7-HCAA cofactor PrP^Sc^ and mice inoculated with WT cofactor PrP^Sc^ **(Supplemental Figure 2)**. Taken together, these data indicate that W99 7-HCAA cofactor PrP^Sc^ is an infectious prion similar to WT cofactor PrP^Sc^.

**Fig 2:**
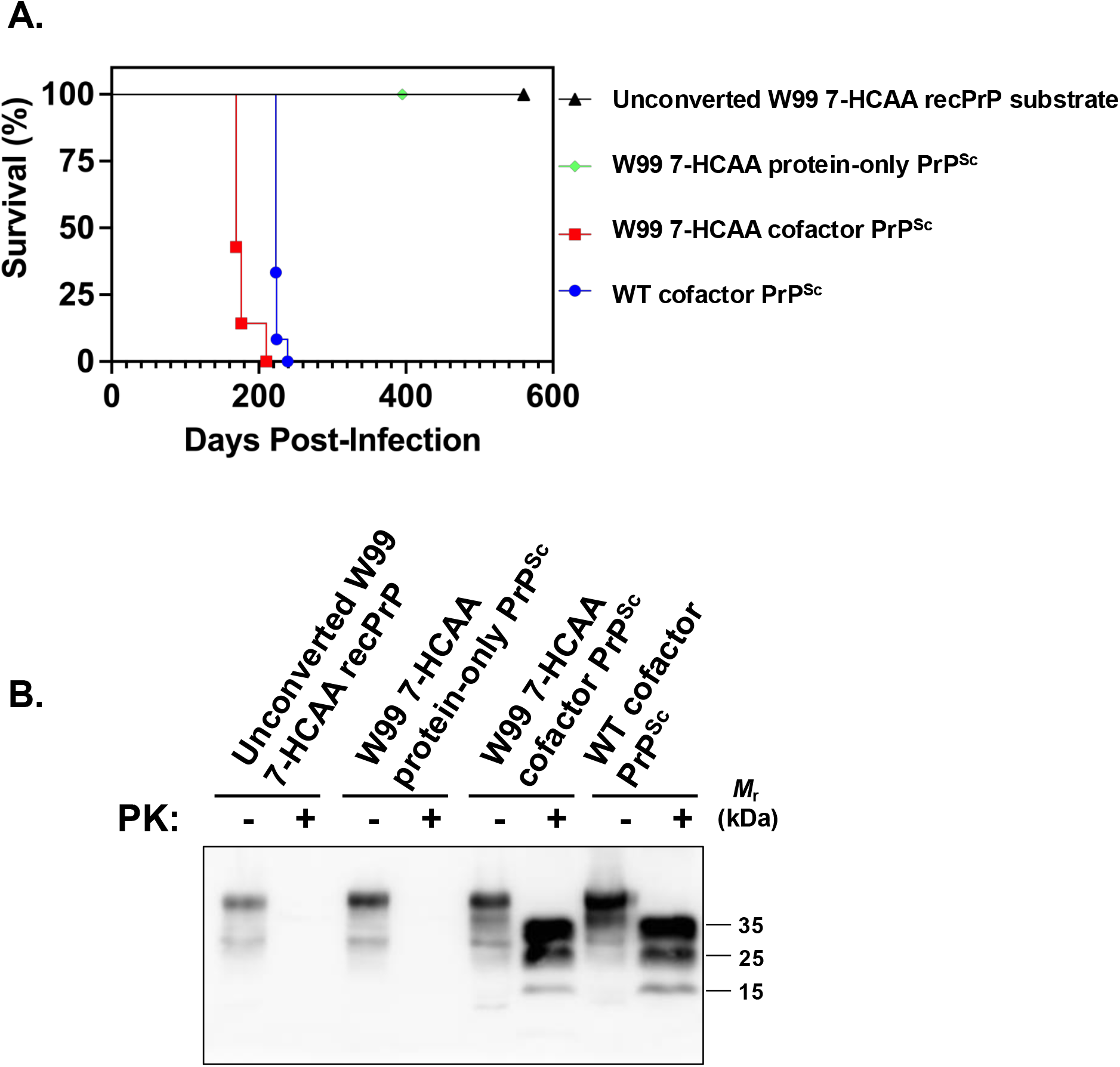
Bioassay in kiBV M109 PrP mice. **(A)** Kaplan-Meier scrapie-free survival plots for mice inoculated intracerebrally with various inocula, as indicated. W99 7-HCAA cofactor PrP^Sc^ n=7; all the other samples: n=12. **(B)** Western blot of brain homogenates from mice inoculated with various samples, as indicated. Samples were digested with proteinase K (PK) where indicated.

**Fig 3:**
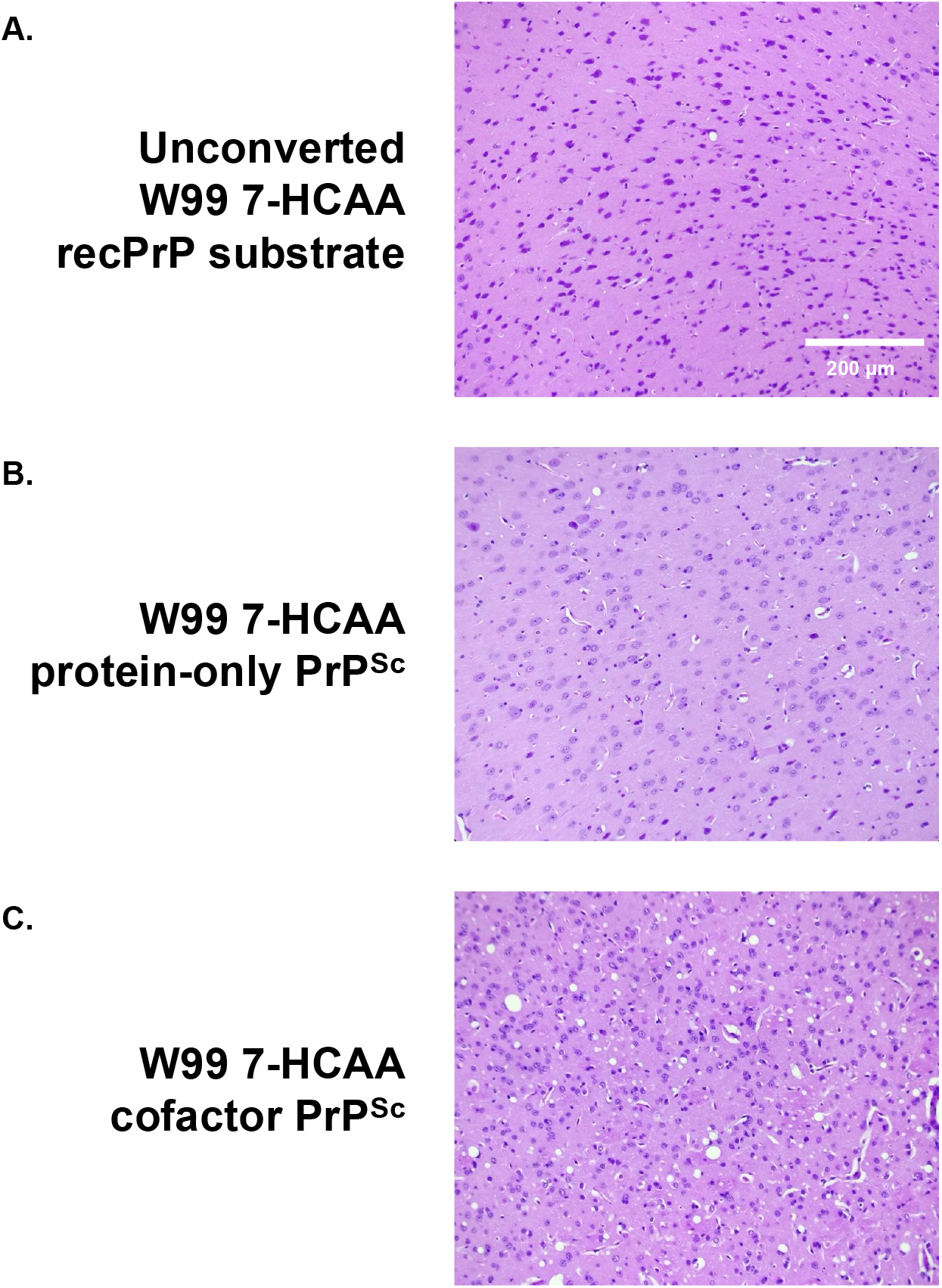
Neuropathology in kiBV M109 PrP mice. Representative hematoxylin and eosin (H&E)-stained microscopic images of cerebral cortex taken from mice inoculated with various inocula, as indicated.

### 7-HCAA fluorescence levels vary due to conformational changes from recPrP to PrP^Sc^

7-HCAA is a non-canonical amino acid whose fluorescence is sensitive to polarity and pH*(24, 25)*. To determine whether 7-HCAA can report on conformational changes during PrP misfolding, we compared the fluorescence emission spectra of W99 7-HCAA PrP in its native (recPrP) and misfolded (PrP^Sc^) states. For each sample, emission spectra were collected under both non-denaturing and denaturing conditions, with the latter serving as an internal standard to estimate maximal fluorescence upon full solvent exposure of the probe.

We first confirmed that the emission spectra of free 7-HCAA dye in non-denaturing and denaturing conditions were indistinguishable, ensuring that spectral differences observed in protein samples arise from conformation-dependent quenching rather than buffer composition **(Figure 4D, overlap between red squares and black circles)**.

**Fig 4:**
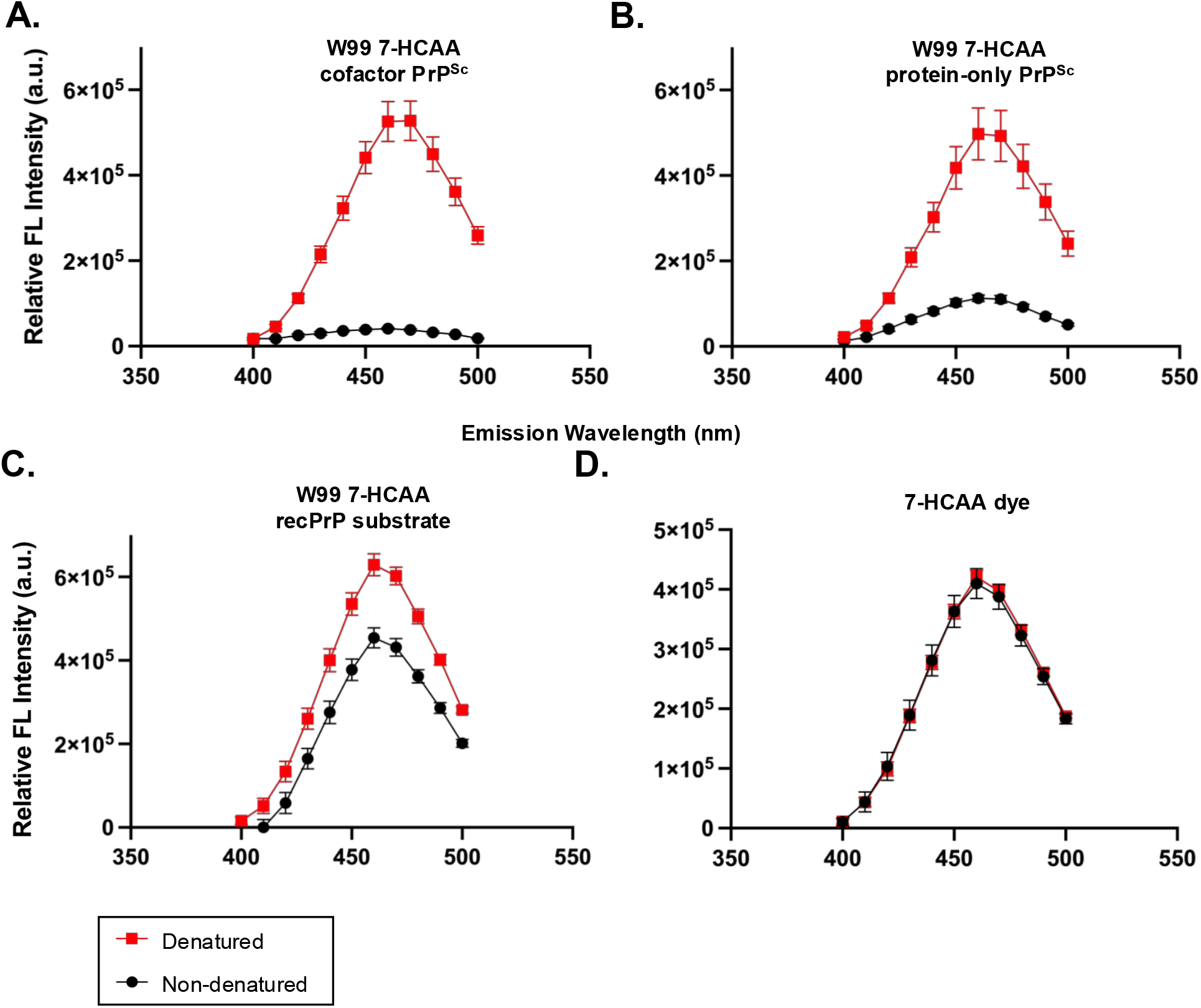
Fluorescence spectra of W99 7-HCAA conformers. Fluorescence emission spectra of various samples following excitation at 360 nm. Red boxes = fluorescence in denaturing conditions; Black circles = fluorescence in non-denaturing conditions (indicates maximal potential fluorescence). **(A)** W99 7-HCAA cofactor PrP^Sc^. **(B)** W99 7-HCAA protein-only PrP^Sc^. **(C)** Unconverted W99 7-HCAA recPrP substrate. **(D)** Free 7-HCAA dye (reagent). Mean values ± SEM are shown (n = 3).

We next analyzed W99 7-HCAA cofactor PrP^Sc^ and W99 7-HCAA protein-only PrP^Sc^ conformers. Under non-denaturing conditions, both samples exhibited relatively low levels of fluorescence intensity **(Figure 4A-B, black circles)**. Upon denaturation, fluorescence intensity at 460 nm increased ∼13-fold for W99 7-HCAA cofactor PrP^Sc^ and ∼4-fold for W99 7-HCAA protein-only PrP^Sc^ **(Figure 4A-B, red squares)**, indicating that W99 is largely buried in the fibrillar state and becomes solvent-exposed upon denaturation. Notably, W99 7-HCAA cofactor PrP^Sc^ displayed a higher denatured/non-denatured fluorescence intensity ratio than W99 7-HCAA protein-only PrP^Sc^ (ratio W99 7-HCAA protein-only PrP^Sc^ versus ratio W99 7-HCAA cofactor PrP^Sc^, two-tailed Welch’s t-test (p = 0.0302)), suggesting differences in local conformation at this site.

We then examined the native W99 7-HCAA recPrP substrate. In non-denaturing conditions, fluorescence intensity was already high, indicating that residue 99 in this soluble conformer is largely exposed to aqueous solvent. Upon denaturation, fluorescence increased only ∼1.3-fold **(Figure 4C, compare red squares and black circles)**.

Together, these results demonstrate that 7-HCAA emission spectra can distinguish between native and misfolded PrP states by reporting on residue-specific solvent accessibility, while denaturation provides a reference for maximal probe exposure.

### Fluorescence of PK-digested samples reveals residue incorporation into the PK-resistant core

To further probe 7-HCAA as a reporter of environment-dependent conformational changes, we designed an assay combining PK digestion with fluorescence readout. PK selectively cleaves and releases solvent-accessible regions, whereas protease-resistant regions remain attached to insoluble amyloid fibrils. Following PK digestion, samples are centrifuged at 18,000 × *g* for 20 min to separate soluble fragments (supernatant) from insoluble PrP^Sc^ fibrils (pellet). All fractions are subsequently denatured prior to fluorescence measurement to expose 7-HCAA residues that would otherwise be quenched when buried within fibrils. Thus, fluorescence intensity reflects the total amount of 7-HCAA-labeled protein present in each fraction.

We applied this assay to W99 7-HCAA recPrP substrate, W99 7-HCAA protein-only PrP^Sc^, and W99 7-HCAA cofactor PrP^Sc^. In the absence of PK, fluorescence is only found in the pellet fraction for both W99 7-HCAA protein-only PrP^Sc^ and W99 7-HCAA cofactor PrP^Sc^ samples **(Figure 5B and 5C, first bar pairs)**. This is expected, as these samples are fully converted into insoluble fibrils that sediment upon centrifugation. In contrast, the W99 7-HCAA recPrP substrate control remains soluble, with fluorescence predominantly in the supernatant **(Figure 5A, first bar pair)**.

**Figure 5:**
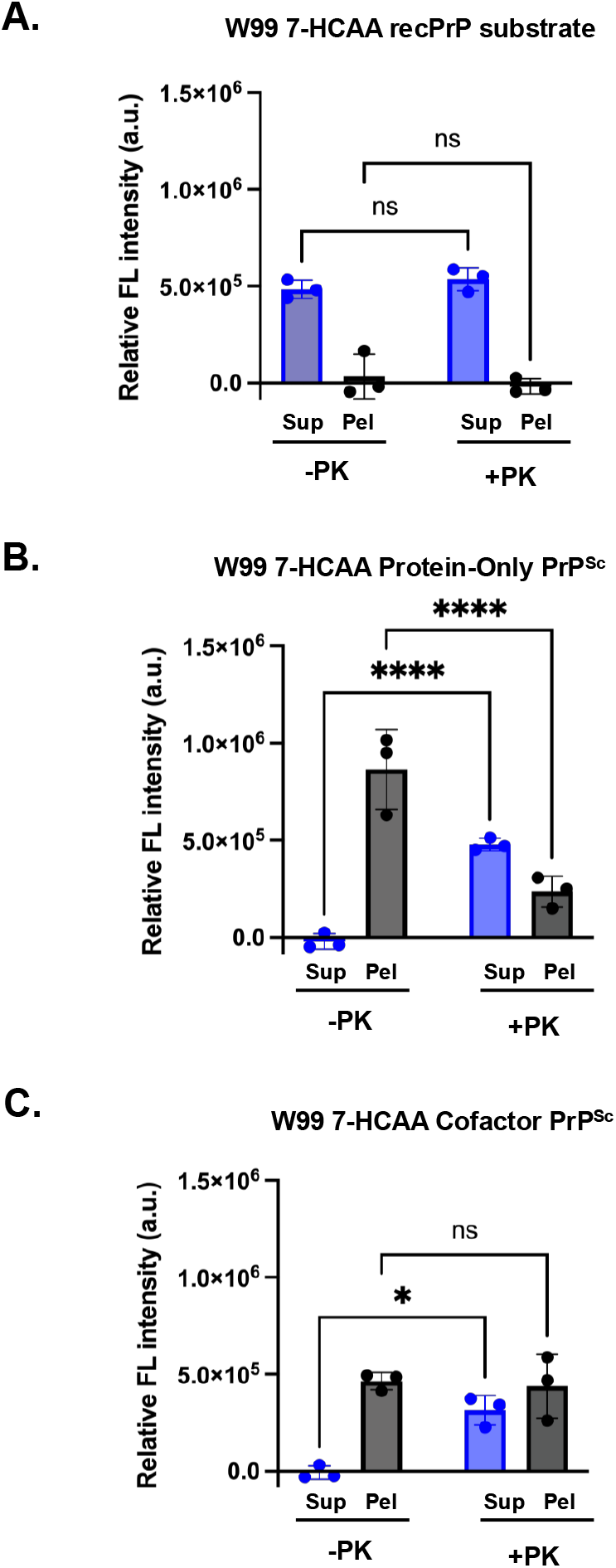
Fluorescence measurements in soluble and insoluble fractions following protease-digestion and centrifugation. Various samples, as indicated, were either treated with PK (+PK) or not (-PK) and centrifuged at 18,000 × g for 20 min to separate into supernatant (Sup) and pellet (Pel) fractions, as indicated. **(A)** W99 7-HCAA recPrP substrate. **(B)** W99 7-HCAA Protein-Only PrP^Sc^. **(C)** W99 7-HCAA Cofactor PrP^Sc^. All reactions were denatured by boiling for 30 min at 60°C in 6 M guanidine HCl, 20 mM Tris, pH 7.4 prior to fluorescence measurement. Mean values ± SEM are shown (n = 3).

Following PK treatment, a redistribution of fluorescence to the supernatant is observed for both PrP^Sc^ conformers **(Figure 5B and 5C, second bar pairs)**. However, not all fluorescence is released: ∼33% of the fluorescence remains in the pellet for protein-only PrP^Sc^, while a larger fraction (∼58%) remains in the pellet for cofactor PrP^Sc^. This is consistent with the presence of protease-sensitive PrP^Sc^ species (sPrP^Sc^), as previously reported*(43, 44)*. As expected, all the fluorescence in the W99 7-HCAA recPrP substrate sample remains in the supernatant after PK treatment, consistent with its fully protease-sensitive nature **(Figure 5A, second bar pair)**.

Overall, this assay enables detection of regions that are hydrophobic but still partially PK-sensitive, providing a simultaneous readout of the local environment of specific residues and their incorporation into the PK-resistant core in various PrP conformers.

## Discussion

We report a fluorescence-based method capable of detecting site-specific local conformational changes during the process of infectious prion formation. By using 7-HCAA PrP molecules as substrates in an *in vitro* system that efficiently generates infectious prions, this platform provides a uniquely powerful approach to probe the structural dynamics of prion conversion.

### A unique platform to study the dynamic process of prion formation

The conversion of PrP^C^ to PrP^Sc^ is a critical step in prion-disease pathogenesis. Although the structures of both conformers are known*(5-10)*, the mechanism of structural conversion remains elusive. It is technically challenging to study this process because PrP^Sc^ amyloid fibrils (and potentially misfolded intermediates) are aggregated and insoluble. Some biophysical techniques such as solution nuclear magnetic resonance (NMR) can be used to study the folding of globular proteins, but are not suitable for insoluble proteins. Other methods, including deuterium-exchange mass spectrometry (DXMS), solid-state NMR, and cryo-EM, can be used to study end-state amyloid fibril structures but cannot detect intermediate states. Computational approaches have been used to model the dynamics of prion conversion*(11, 12)*, but an experimental method is needed to test such predictions.

Substitution with 7-HCAA has previously been used to probe conformational changes in globular proteins*(21, 24, 45, 46)*. However, this strategy is particularly well-suited for studying the process of infectious prion formation, a sensitive conformational process that is easily disrupted by relatively small perturbations. 7-HCAA is similar in size and structure to aromatic amino acids, so substitution is relatively conservative, which is important for maintaining PrP^C^ conformation and ability to form infectious PrP^Sc^. By contrast, conventional labeling strategies use bulky chemically reactive reagents that modify proteins non-specifically at multiple sites*(47)* and therefore cannot be used with PrP^C^ substrates, which are sensitive to structural perturbations. Another important advantage of our approach is that we are able to selectively replace individual residues with 7-HCAA*(23, 36)*.

The application of 7-HCAA to study prion formation specifically exploits the sensitivity of the fluorophore to environmental polarity. In the predominantly α-helical PrP^C^ state, 7-HCAA substituted at most residues would be expected to produce strong fluorescence due to a high degree of solvent accessibility, as previously determined by DXMS*(48)*. As misfolding proceeds and the labeled residue becomes buried within the β-sheet-rich amyloid fibril, the local environment becomes more hydrophobic, likely causing 7-HCAA fluorescence to quench. This conformation-dependent reduction in fluorescence intensity provides a rapid, sensitive, and quantitative measure of misfolding for the PrP domain containing 7-HCAA.

A unique feature of our *in vitro* conversion system that is well-suited for fluorescence analysis is its relatively high conversion efficiency*(32-35)*, which optimizes the signal-to-noise ratio. Another important feature of our *in vitro* system is that it can produce infectious prions with high levels of specific infectivity*(41, 49)*. Thus, our platform can be used to investigate a structural transition process that is relevant to understanding how infectious prions are formed.

### Applications and limitations

We envision that this novel platform can be used to determine the order in which different PrP^C^ domains are sequentially incorporated into the PrP^Sc^ structure during the process of infectious prion formation. A panel of PrP substrates with 7-HCAA substituted into different domains could be used to determine the relative kinetics of destabilization and conformational change for each domain during the conversion process.

A limitation of our work is that we have thus far only purified and tested a single 7-HCAA substrate (W99 7-HCAA recPrP) for its ability to convert into PrP^Sc^. Substitution of 7-HCAA for different residues could destabilize PrP^C^ structure and/or disrupt its ability to convert into PrP^Sc^. This potential problem may be more likely for substitution of non-aromatic residues. However, since every domain contains multiple residues, it is likely that some substitutions will be tolerated in each domain.

Another limitation of this platform is that a single 7-HCAA substitution cannot be used to distinguish whether a change in fluorescence intensity reflects local side-chain repacking or a large-scale domain rearrangement. One way to distinguish between these different scenarios would be to integrate and model data from multiple substrates with 7-HCAA substituted at different positions within the same domain. Another possibility would be to pair the genetically encoded fluorophore with additional labels for Förster resonance energy transfer (FRET)*(50)*. FRET measurements would report on long-range intramolecular distances and could reveal contacts between specific domains during misfolding, as well as potential interactions between PrP and cofactor molecules. Because 7-HCAA can serve as a FRET donor*(51-53)*, and its position can be varied across the protein, this approach would enable systematic mapping of distance changes throughout the conversion process, information that is currently inaccessible with existing methods.

Interestingly, our data show that 7-HCAA fluorescence can also be used to probe site-specific structural differences between different PrP^Sc^ conformers. Specifically, we observed that W99-7-HCAA cofactor PrP^Sc^ exhibits lower fluorescence intensity than W99-7-HCAA protein-only PrP^Sc^. This suggests that residue 99 is buried in a less polar environment within the cofactor PrP^Sc^ structure than within the protein-only PrP^Sc^ structure. This observation is consistent with prior DXMS work showing that the region spanning residues 91–115 is more solvent-accessible in protein-only PrP^Sc^ than in cofactor PrP^Sc^*(48)*.

Beyond its use here to study infectious prions, 7-HCAA and other environmentally sensitive ncAAs could also be used to study the conversion mechanism of other amyloidogenic proteins. Disease-associated proteins such as Aβ, tau, α-synuclein, and SOD1 each transition from soluble states to insoluble, β-sheet-rich amyloid fibrils, yet the structural sequence of these transitions is largely unknown because of technical limitations similar to those that have previously confounded the study of PrP misfolding*(54-57)*. Placing an environmentally sensitive fluorophore at defined positions in any of these other proteins would enable real-time, domain-level mapping of conformational changes during the amyloid formation process, information that non-specific amyloid dyes such as Thioflavin-T cannot provide*(58)*. A similar approach can also be used to study the conformational change mechanisms of functional amyloids such as curli fibers, CPEB aggregates, or Pmel17 fibrils, where controlled polymerization is essential for biological function*(59-61)*.

In summary, we report a novel system that couples site-specific incorporation of a fluorescent, environmentally sensitive ncAA into PrP with a high-efficiency *in vitro* prion conversion assay. This approach opens the possibility of monitoring local structural changes during the dynamic process of infectious prion formation and has broad applicability to other amyloids.

## Accession IDs

M109 BV PrP 23-231 (UniProt ID: Q8VHV5_MYOGA)

