## Supplemental Figure for "Fluorescent non-canonical amino acid as a site-specific conformational probe of prion formation"

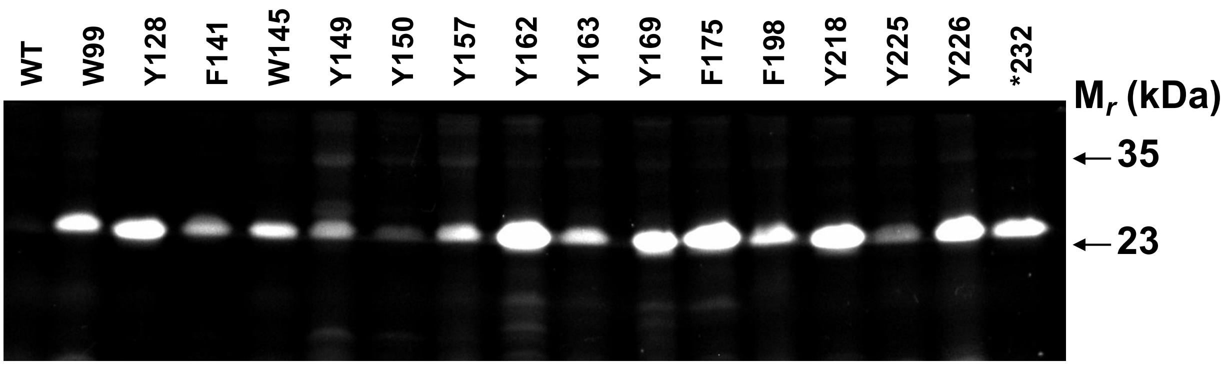


***Supplemental Figure 1: Expression of various 7-HCAA mutant PrP molecules in E. coli.*** *Fluorescence-imaged SDS polyacrylamide gel of crude lysates prepared from bacteria expressing mutant PrP molecules with 7-HCAA substituted for various residues, as indicated. WT = control bacteria expressing wild-type PrP without 7-HCAA substitution. *232 = amber codon was inserted immediately after the final residue S232 to produce full-length PrP with 7-HCAA added to the C-terminus.*

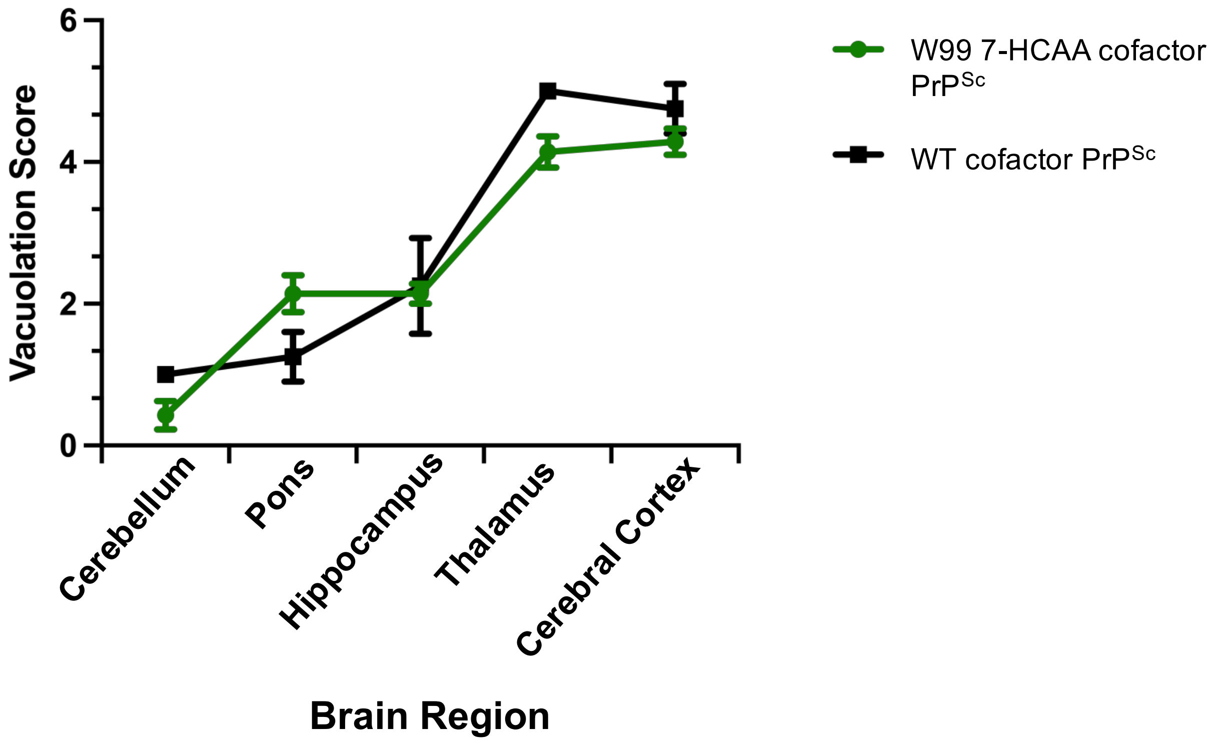


***Supplemental Figure 2: Regional neuropathology of kiBV M109 PrP mice inoculated with W99 7-HCAA cofactor PrP$^{Sc}$ and WT cofactor PrP$^{Sc}$***. *Profiles of vacuolation scores of animals inoculated with either W99 7-HCAA cofactor PrP$^{Sc}$ (green circles) or WT cofactor PrP$^{Sc}$ (black squares). Mean values ± SEM are shown.*
